# Optimizing Design of Genomics Studies for Clonal Evolution Analysis

**DOI:** 10.1101/2024.03.14.585055

**Authors:** Arjun Srivatsa, Russell Schwartz

## Abstract

Genomic biotechnologies have seen rapid development over the past two decades, allowing for both the inference and modification of genetic and epigenetic information at the single cell level. While these tools present enormous potential for basic research, diagnostics, and treatment, they also raise difficult issues of how to design research studies to deploy these tools most effectively. In designing a study at the population or individual level, a researcher might combine several different sequencing modalities and sampling protocols, each with different utility, costs, and other tradeoffs. The central problem this paper attempts to address is then how one might create an optimal study design for a genomic analysis, with particular focus on studies involving somatic variation, typically for applications in cancer genomics. We pose the study design problem as a stochastic constrained nonlinear optimization problem and introduce a simulation-centered optimization procedure that iteratively optimizes the objective function using surrogate modeling combined with pattern and gradient search. Finally, we demonstrate the use of our procedure on diverse test cases to derive resource and study design allocations optimized for various objectives for the study of somatic cell populations.

## 1 Introduction

Recent developments in genomic biotechnology have allowed for increasingly precise characterization and manipulation of genomic, epigenomic, and transcriptomic content at a single-cell level (Li *et al*., 2021). Our growing understanding of somatic variability has paved the way for understanding biological processes – like aging and development – and also disease conditions, like cancer (Olafsson and Anderson, 2021). Though genomic medicine is still in its infancy, the future might involve various genome monitoring and engineering strategies in the management of these biological processes (Das *et al*., 2015).

The realm of cancer biology has proven to be especially fruitful for biotechnological and genetic analyses. Cancer is a heterogeneous disease whereby a process of somatic evolution leads to an accumulation of genetic modifications that gradually give rise to clonal disorder and neoplasias. Cancer cells are typically imbued with one or more of many possible hypermutation phenotypes that produce characteristic patterns of genetic or epigenetic plasticity in their descendants(Loeb, 2001). Examples include the chromosome instability (CIN) phenotypes characteristic of TP53 defects (Pino and Chung, 2010), point mutation phenotypes of DNA polymerase defects (Barbari and Shcherbakova, 2017), or various phenotypes of DNA mismatch repair defects (Peltomäki, 2001). Furthermore, neoplasms often contain complex scaffolds of putatively healthy and neoplastic cells, each exhibiting varying genetic and epigenetic changes and spatial distributions within the tumor microenvironment (Martincorena *et al*., 2017). The clonal evolution from a healthy cell population to an invasive, metastatic one is still an unsolved problem – a primary challenge being the wide variability present within each neoplasm. As such, the ushering of personalized care, where an individual patients’ neoplasias might be analyzed for mutability patterns and targeted strategically, holds great promise.

Advancing personalized cancer care will hinge in part on our ability to develop rigorous statistical and data science frameworks for dissecting the complexity of the biology of somatic mutability, involving decision making in the face of many complementary genomic technologies now available to us. Current statistical and algorithmic approaches take various genetic readouts as inputs and give insights into the population structure and dynamics of cell populations(Ding *et al*., 2014). Population features we might want to characterize include a varying and expanding set of characteristics including: cellular phylogenetic trees, mutations (structural, point, and copy number changes) and mutability patterns (mutational signatures), spatial distributions, epigenetic alterations, and transcriptional activity (Grody *et al*., 2023). Researchers seeking to characterize such features have available to them numerous various combinations of genomic tools combined with a wide variety of algorithms with which to draw inferences (Pareek *et al*., 2011; Schwartz and Schäffer, 2017). We now have scalable (bio)technological tools and associated algorithms allowing us to measure changes in RNA, DNA, translation, transcription, and the epigenome. We also have tools that allow us to modify DNA/RNA/epigenome information at the single cell level allowing for more involved and accurate information collection systems (e.g., CRISPR barcoding). This wealth of options is making ever more complicated the key question of how a researcher can design a maximally efficacious study to gather specific information about a somatic evolution process within resource constraints. This problem is especially difficult because of the high dimensional study-design optimization space combined with arbitrarily-complex cell populations without ground-truth data. Multiomic designs (e.g., combining various sequencing modalities) may be particularly valuable for information gain but present a challenge for traditional optimization algorithms (Mangiante *et al*., 2023). Working effectively and efficiently with the complex space of options requires principled alternatives to the current practice of essentially *ad hoc* design.

In this paper, we aim to meet this challenge by developing tools for design of studies of somatic evolution processes using ideas from Bayesian optimization. In doing so, we define a generalized study design allocation problem for cancer genetics studies under experimental, simulation, and budget constraints. We then show that our optimization procedure can find effective study designs for diverse somatic mutability analyses via a set of case studies.

## 2 Materials and Methods

In a typical cancer use-case, a researcher might be interested in any subset of the genetic, epigenetic, transcriptional, or proteomic information contained within a cell population of interest. In order to answer the general study-design problem, we first pose a formal definition of a study design along with a statement of the optimization problem. The following section defines the variables and mathematical framework in detail. The main components of the study design problem are represented by bold text. Somatic evolution instances and their associated genetic study designs are generated by the simulator in Srivatsa *et al*. (2023).

### 2.1 The Optimization Problem

Our definition requires the following components

1. In this paper, we define a **study design, x**, as a vector:

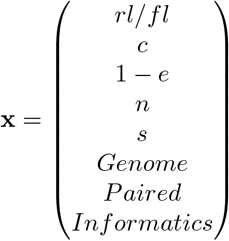

Detailed definitions of each of these variables is found in Appendix Table 4; here each of the sequencing variables are condensed into some representative value for the study design.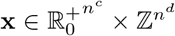, where *n*^*c*^ denotes the number of continuous variables and *n*^*d*^ defines the number discrete variables. Many of these variables, like coverage or read length, might be considered discrete, but have a continuous interpretation. Discrete variables may be integer values (e.g., read length) or categorical (e.g. variant caller).
2. We next define a **budget function**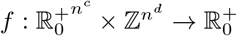 and a **cost function**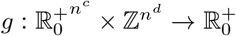 Informally, we think of the budget function *f* as assigning a monetary cost to a particular study design and the cost function *g* as assigning a fraction of the maximal budget used by a particular study design. We assume that these two functions are quickly computable for any design **x**. However, the functions may lack a closed form expression and may not be continuous or differentiable.
3. **Constraints** for the design are given by *f* (**x**) ≤ *b*, where *b* denotes a maximum budget. One might define these constraints by partitioning the domain into continuous and discrete sets and defining valid subsets for each, though any other function could be used.
4. Central to the optimization framework is our **loss function** *L*_*q*_(**x**). *L*_*q*_ :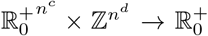 measures the error with which our study design returns a particular property or set of properties the somatic system studied. There is a dependency on the biology of the somatic cells themselves, described by the *q* subscript in the loss function, where a cell population is generated from a stochastic realization *q* ∼ *Q*(*t*). This loss function is typically difficult to compute in a cancer study. Hence, there may only be a limited number of real and simulated loss function evaluations that can be used in our optimization solution.
5. A **regularization penalty** *λ* ∈ ℝ^+^ balances the efficacy of a study design and its resource cost.
6. The **score** of a study design is defined as *λg*(**x**) + *L*_*q*_(**x**), which measures the overall efficacy of a target study design **x**. Note that in our framework, lower scores imply more effective study designs.

We are then prepared to offer our formal problem statement:

**The Study Design Problem** Given a fixed simulation or experimental budget to iteratively generate *M* datasets, 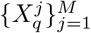 of the form 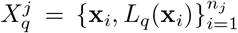 corresponding to study design evaluations on a somatic evolution process, we seek to solve the following constrained optimization problem:

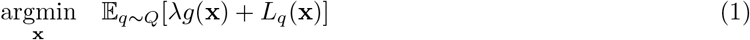

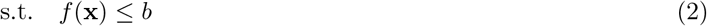

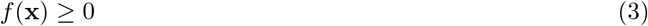

Our formulation amounts to a stochastic black-box mixed-integer programming problem. The problem is additionally challenging because our loss function is potentially discontinuous and costly to estimate, and can only be sampled in batches via simulation. Various approaches have been used in the optimization literature to solve similar problems (Liuzzi *et al*., 2012; Kelley, 2002). Some rely on the construction of surrogate functions which are used as either first-pass solutions or aids in traditional iterative search methods (Gramacy, 2020). Others estimate gradients directly from simulation calls and use these to approach minima iteratively (Spall, 2012). Yet others do not rely on gradients at all, instead employing direct search style algorithms that iteratively explore promising points (Audet and Dennis Jr, 2001). Depending on assumptions regarding our function and input domain, some algorithms have proveable convergence guarantees while others remain heuristics(Sriver, 2004). More recent methods have expanded such strategies to mixed variable domain sets. For reviews of various algorithms for related problems see Abramson (2003); Sriver (2004). The idea of using sequentially simulated somatic evolution instances in combination with diverse biotechnology study design instances is, to our knowledge, new to the cancer genomics literature and there is thus a dearth of methods for the study design problem. Software for more generic optimization is poorly suited to the particulars of this domain, failing to encapsulate the stochastic and sequential nature of our sampled points, the mixed variable domain of our study design vector, or parallelized simulation and compute scheme. We thus develop a new approach, borrowing effective ideas from past literature, such as mesh-search for mixed variable programming and surrogate modeling, in designing an effective and efficient optimization method.

### 2.2 Optimization Strategy

Our overall optimization strategy uses an iterative exploration-exploitation approach to return progressively better solutions to the study design problem. Given the high dimension of the search space and cost per experiment, we balance exploration of high variance regions of the search space with local exploration of already promising points. The initial stage of the algorithm produces a sampling matrix of study designs for both the cost function and loss function, which are then used to build initial surrogate models for the combined cost and loss function. The algorithm proceeds by using the surrogate functions to simultaneously explore high uncertainty regions of the sample space and perform local neighborhood search of current high performance study designs. After each iteration, the surrogate functions are updated, and a queue of high performance points is iterated based on our latest local and global search. During the algorithm, cost constraints are maintained by rejecting proposed designs that are over the allowed budget. The algorithm ends when either the simulation budget is exhausted or it fails to find a significantly better point after a fixed number of iterations. Figure 1 provides a flowchart of the algorithm, Full pseudocode is found in Algorithm 1 in Appendix section 6.2.

**Fig. 1.**
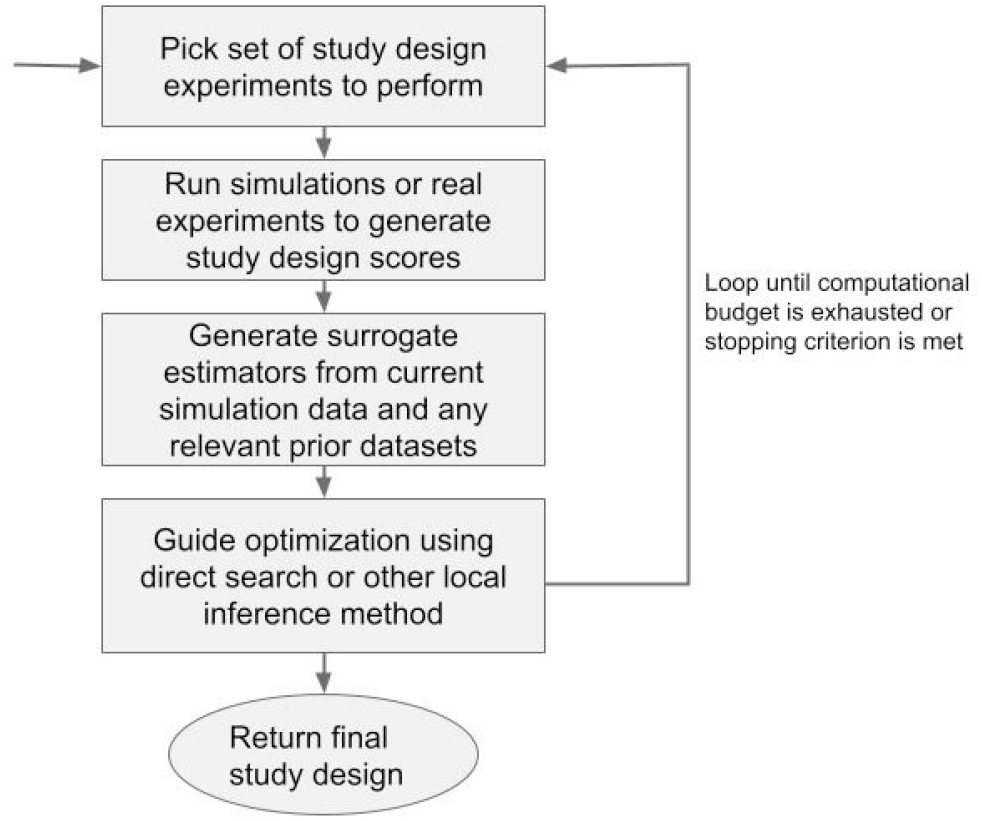
A flowchart of the main algorithm is depicted above. The algorithm functions by repeatedly estimating a surrogate functions from simulation calls and exploring promising regions of the search space.

#### Generating Experimental Designs

At each iteration, the algorithm must estimate the loss function and potentially cost function. Here, we assume that we can pick the subsequent set of study designs via simulation. During the initial stage of the algorithm, we do not have any information about which study designs are most effective, and therefore want a set of study designs that “fills” the domain space. We use Latin hypercube sampling (LHS)(McKay *et al*., 2000) for this purpose. Instead of simply sampling from the joint distribution, we partition each study design variable **x**^*i*^ (the superscript is used to differentiate between the variable and the data point number) into *n*_0_ equally-sized intervals spanning the range of the variable (where *n*_0_ is the number of samples). We randomly sample from each interval to generate a set of *n*_0_ samples for variable **x**^*i*^. This is repeated for each variable **x**^*i*^, and samples from each variable are concatenated in a random permutation for each variable to generate a full matrix 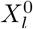 of dimension *n*_0_ × *d*. Here *d* represents the full dimension of a single vector study design **x**, i.e. *d* = *n*^*c*^ + *n*^*d*^ the total number of discrete and continuous variables. The set of samples produced by LHS in promotes variability in the sample space, however the sample space is still exponentially large. We use the stochastic evolutionary algorithm of Jin *et al*. (2003) to generate a favorable LHS sample via a maximin distance i.e., max 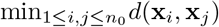 where *d*(**x**_*i*_, **x**_*j*_) denotes the (Euclidean) distance between samples. This LHS sample is intended to maximize the minimal distance between any two sample rows in the data matrix – thereby ensuring a more even spread in the data. Since some of these designs may exceed the cost function budget, we reject these samples and generate LHS matrices until the target sample size is exceeded. If the cost function is not obtainable in closed form, we might also want to approximate this function, and would similarly draw samples for a matrix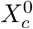. Data matrices 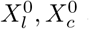 are passed to the next stage of our algorithm.

#### Generating Surrogate Models

Using our initial set of data 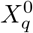, we then set out to develop a Gaussian process estimator for our loss function 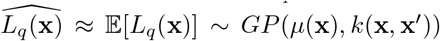. The Gaussian process is defined by a mean and covariance function estimating similarity between points in the input space (and encouraging smoothness). Here we use the Matern 3/2 covariance function defined below.

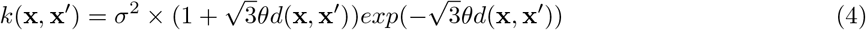

In the above kernel, *d* represents some distance metric and *θ* represents a smoothness hyperparameter. The mean and variance of new sample points can be estimated using a conditional distribution derived from the definition of the Gaussian process (c.f., Rasmussen (2003)), i.e. :

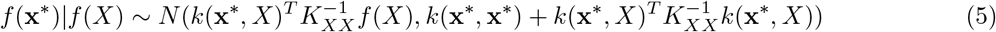

Here *X* denotes the set of already observed study design vectors, *K*_*XX*_ denotes the pairwise kernel matrix of our *X* vectors, and *k*(**x**^*^, *X*) denotes a column vector of kernels of **x**^*^ with each element in *X*. Traditional Gaussian process models can be extended to mixed variables by adapting the kernel distance function (Saves *et al*., 2023a). Generally the kernel function is factored into continuous and discrete kernels, with the overall kernel/similarity defined as their product. We use the exponential homoscedastic hypersphere kernel for categorical variables (c.f., Saves *et al*. (2023a),Garrido-Merchán and Hernández-Lobato (2020)). These functions and inferences are implemented in the Surrogate Modeling Toolbox package (Saves *et al*., 2023b).

If we do not have a closed form expression for our cost function *g*(**x**), we can also generate an estimator 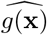 using our dataset 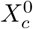. We use our Gaussian surrogate function(s) to rapidly compute our overall score function for various study designs: 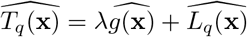

### Exploring Additional Points

A critical stage of the overall optimization strategy is the selection of a new set of study design experiments to generate labeled loss function scores for given previous sets of data 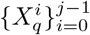. There are three main methods we use to select points: Gaussian process acquisition function selection, gradient descent iteration, and mesh search iteration. Given *n*_*j*_ available sampling points for iteration *j*, we define an exploration coefficient, *e*_*j*_, and generate a large number of points via an LHS strategy. The LHS sample is winnowed via the lower confidence bound acquisition criteria for Gaussian processes. We select the minimum ⌊*e*_*j*_*n*_*j*_ ⌋ points of the LHS sample to generate for the next iteration via the values of the LCB function, which is defined by the function below:

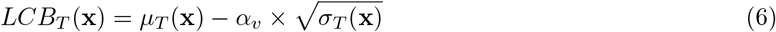

*α* represents an exploration parameter which emphasizes high variance points. This strategy ensures that we explore “promising” points with low values and/or high variance given our current set of explored points.

The algorithm also explores in the vicinity of the current best simulated points via local search in an “exploitation” phase. At the current iteration, the other *n*_*j*_ − ⌊*e*_*j*_*n*_*j*_⌋ points are explored via local search with *n*_*m*_ dictating the number of points explored via mesh search and *n*_*g*_ the number of points explored via gradient descent. The current minima is designated a “mesh center”, around which a mixed-integer mesh is built to sample. The minimal point can be partitioned into continuous and discrete components, **x** = (**x**^*c*^, **x**^*d*^). A mesh is defined by a set of directions, *D*(**x**, *j*), and a “fineness” parameter *Δ. D*(**x**, *j*) is a matrix of dimension *n*_*c*_ × |*D*|, with the columns defining a positive spanning set of the continuous space, *Θ*^*c*^(informally a set of directions to continue the continuous search). The mesh is defined as:

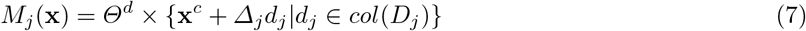

Informally, we can picture the mesh as a set of lattices surrounding the input point. Each discrete value set, input point, and iteration may have a different lattice. Given we have a fixed simulation budget, we cannot evaluate the entire mesh and instead probabilistically pick from the continuous neighborhood lattice of the current point, the discrete neighbors of the current point, and, potentially, the continuous neighborhood of the discrete neighbors – with each set defined below. :

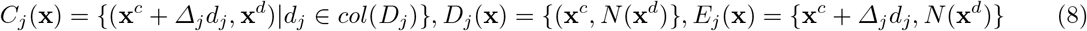

We first randomly sample points in the continuous neighborhood lattice (*C*_*j*_(**x**)) of our current mesh center. We then progress to a discrete neighborhood (*D*_*j*_(**x**)) of our current point; for categorical variables one might define such a neighborhood by setting each variable equidistant to one another, while for integer variables (e.g., read length) one might set a fraction of the range as a “discrete ball” surrounding a point. With sufficient budget on the current iteration, we explore the extended continuous neighborhoods of our discrete neighbors (*E*_*j*_(**x**)). With a large budget one could exhaust all points in the mesh *M*_*j*_(**x**).

At the remainder *n*_*g*_ = *n*_*j*_ − ⌊*e*_*j*_*n*_*j*_⌋ − *n*_*m*_ of the lowest value points, we perform a gradient-descent search by estimating the gradient numerically at each of the *n*_*g*_ points the surrogate functions. The next point sampled is:

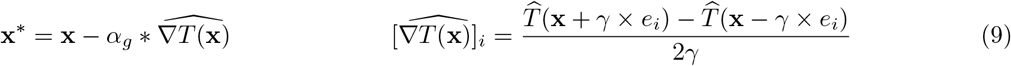

Integer constants *γ* and *α* represent a range of perturbation and movement of the gradient estimator and step (typically both are relatively small). The *e*_*i*_ represents a perturbation vector with a small fraction of the domain range of variable *i* at the *i*^*th*^ position.

In each of the sampling cases, care must be taken to ensure new points are feasible. In the case of points sampled from our acquisition function, we reject points that violate cost constraints. Within our mesh search, we do not allow for the sampling of points outside our domain hypercube. If mesh search points violate our budget, we shrink the fineness of the mesh in hopes of finding a local neighborhood where points are feasible. A similar procedure is implemented in each gradient descent search step, and we also maintain perturbation parameters such that discrete variables remain integer values.

#### Iteration, Stopping Criterion, and Parallelization

The full optimization algorithm involves iteratively picking new data points to evaluate, evaluating sets of data via simulation, and conducting surrogate reestimation of the target function. At any point after the first iteration, a full list of ranked study designs as well as the currently trained surrogate function can be returned to the user. The iteration repeats, exploring high variance points of the surrogate domain while simultaneously exploring points of low expected value until the given budget is exhausted. At each iteration, the exploration parameter and iterated search parameter can be updated (c.f., Appendix Table A1). We generally taper the gradient and mesh search parameters as well as the exploration coefficient with each optimization round.

## 3 Performance Considerations

The main factor in run time of the optimization algorithm is the evaluation of the input points via large scale simulation or experimentation, which incurs a large fiscal, computational, and/or temporal cost. Denoting *S* as the time per simulation batch, our overall run time is therefore proportional to *O*(*MS*), where *M* denotes our simulation batch budget. The simulation experiments and analysis scripts are readily paralellizable across computer processing cores and compute nodes, such that the actual run time is proportional in 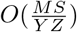 where *Y* and *Z* denote the number of cores and nodes respectively. Storage and maximum memory load per machine must also be considered. Due to the large volume of simulated read data generated, our program deletes the genomic data produced after each batch. Thus, the program requires *O*(*n*_*m*_*G*_*m*_*c*_*m*_*n*_*j*_) units of disk space where *n*_*m*_ denotes the maximal number of single cells in all samples, *G*_*m*_ denotes the max genome size, *c*_*m*_ denotes the maximum coverage, and *n*_*j*_ is the number of samples per iteration. Given that these simulations are resource intensive, our implementation is designed to maximally parallelize the computation while maintaining the memory capacity for each core/node and the storage capacity of the overall cluster. Finally, we note that the simulations themselves should mimic the target biology of the study design problem; one can achieve this by modifying the parameters of the simulation or through data collection strategies.

## 4 Experimental Results

The following section demonstrates the proposed method by posing and solving various hypothetical study design queries. A full study design query is defined by its study design, biology, cost and budget, and loss function parameters. We summarize each use case as it is presented but provide full details in Table 1. The best scores for each experiment are presented in Table 2. Key results are summarized for all experiments in Figs 2-5.

**Table 1.**
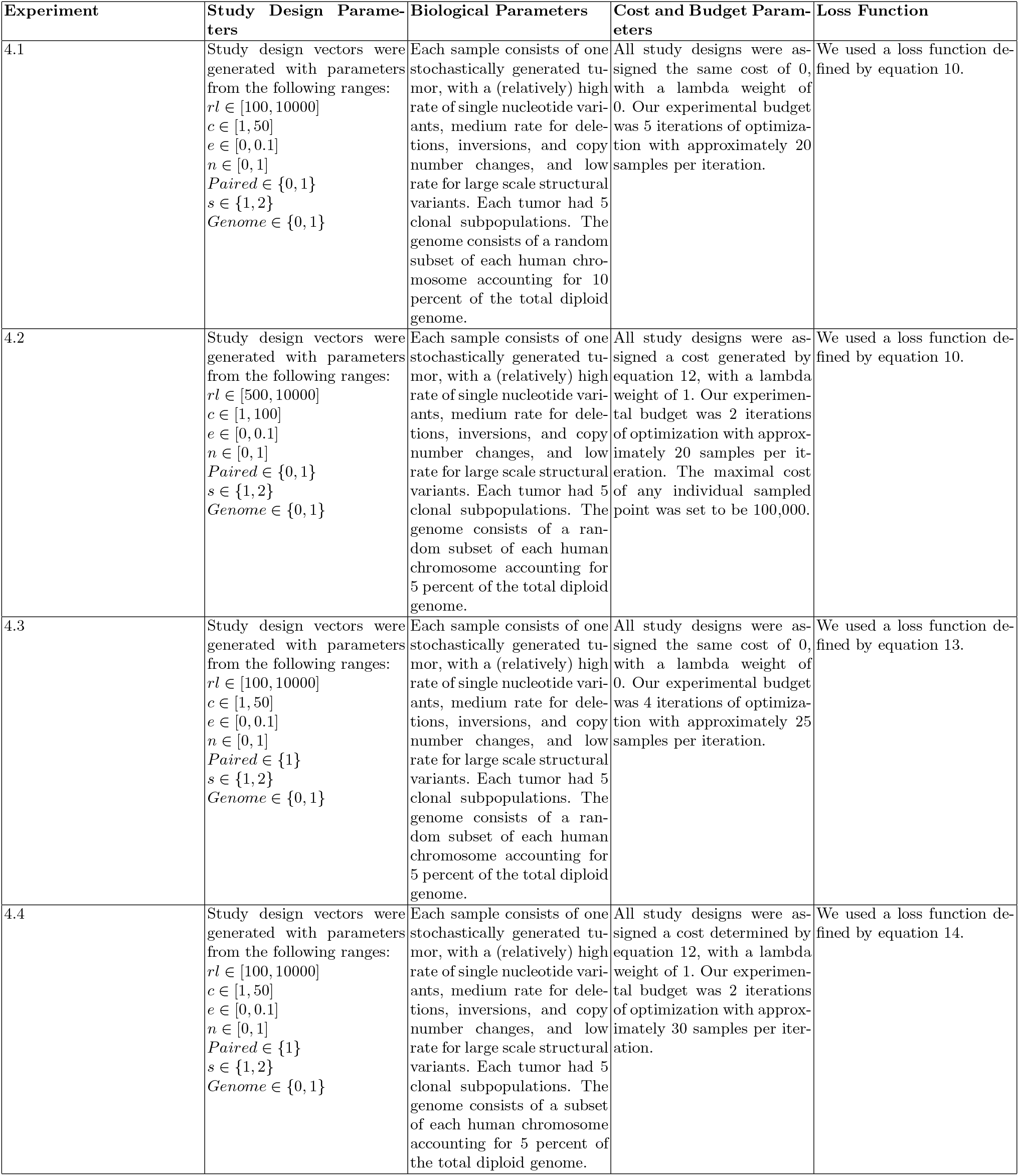
Experimental query settings - including biological, study design, cost function, and loss function parameters-are described below for each performed experiment.

| Experiment | Study Design Parameters | Biological Parameters | Cost and Budget Parameters | Loss Function |
| --- | --- | --- | --- | --- |
| 4.1 | Study design vectors were generated with parameters from the following ranges:<br>$rl \in [100, 10000]$<br>$c \in [1, 50]$<br>$e \in [0, 0.1]$<br>$n \in [0, 1]$<br>$Paired \in \{0, 1\}$<br>$s \in \{1, 2\}$<br>$Genome \in \{0, 1\}$ | Each sample consists of one stochastically generated tumor, with a (relatively) high rate of single nucleotide variants, medium rate for deletions, inversions, and copy number changes, and low rate for large scale structural variants. Each tumor had 5 clonal subpopulations. The genome consists of a random subset of each human chromosome accounting for 10 percent of the total diploid genome. | All study designs were assigned the same cost of 0, with a lambda weight of 0. Our experimental budget was 5 iterations of optimization with approximately 20 samples per iteration. | We used a loss function defined by equation 10. |
| 4.2 | Study design vectors were generated with parameters from the following ranges:<br>$rl \in [500, 10000]$<br>$c \in [1, 100]$<br>$e \in [0, 0.1]$<br>$n \in [0, 1]$<br>$Paired \in \{0, 1\}$<br>$s \in \{1, 2\}$<br>$Genome \in \{0, 1\}$ | Each sample consists of one stochastically generated tumor, with a (relatively) high rate of single nucleotide variants, medium rate for deletions, inversions, and copy number changes, and low rate for large scale structural variants. Each tumor had 5 clonal subpopulations. The genome consists of a random subset of each human chromosome accounting for 5 percent of the total diploid genome. | All study designs were assigned a cost generated by equation 12, with a lambda weight of 1. Our experimental budget was 2 iterations of optimization with approximately 20 samples per iteration. The maximal cost of any individual sampled point was set to be 100,000. | We used a loss function defined by equation 10. |
| 4.3 | Study design vectors were generated with parameters from the following ranges:<br>$rl \in [100, 10000]$<br>$c \in [1, 50]$<br>$e \in [0, 0.1]$<br>$n \in [0, 1]$<br>$Paired \in \{1\}$<br>$s \in \{1, 2\}$<br>$Genome \in \{0, 1\}$ | Each sample consists of one stochastically generated tumor, with a (relatively) high rate of single nucleotide variants, medium rate for deletions, inversions, and copy number changes, and low rate for large scale structural variants. Each tumor had 5 clonal subpopulations. The genome consists of a random subset of each human chromosome accounting for 5 percent of the total diploid genome. | All study designs were assigned the same cost of 0, with a lambda weight of 0. Our experimental budget was 4 iterations of optimization with approximately 25 samples per iteration. | We used a loss function defined by equation 13. |
| 4.4 | Study design vectors were generated with parameters from the following ranges:<br>$rl \in [100, 10000]$<br>$c \in [1, 50]$<br>$e \in [0, 0.1]$<br>$n \in [0, 1]$<br>$Paired \in \{1\}$<br>$s \in \{1, 2\}$<br>$Genome \in \{0, 1\}$ | Each sample consists of one stochastically generated tumor, with a (relatively) high rate of single nucleotide variants, medium rate for deletions, inversions, and copy number changes, and low rate for large scale structural variants. Each tumor had 5 clonal subpopulations. The genome consists of a subset of each human chromosome accounting for 5 percent of the total diploid genome. | All study designs were assigned a cost determined by equation 12, with a lambda weight of 1. Our experimental budget was 2 iterations of optimization with approximately 30 samples per iteration. | We used a loss function defined by equation 14. |

**Table 2.** The best(lowest) scoring study designs for each of the experimental queries is depicted in the table above.

| Experiment | *rl* | *c* | *e* | *n* | *Paired* | *s* | *Genome* | Score |
| --- | --- | --- | --- | --- | --- | --- | --- | --- |
| 4.1 | 4904 | 44.96 | 0.051 | 1 | 1 | 2 | 1 | 0.485 |
|  | 7471 | 43.86 | 0.0 | 1 | 1 | 2 | 1 | 0.492 |
|  | 6023 | 50.0 | 0.079 | 1 | 1 | 2 | 1 | 0.492 |
|  | 6022 | 20.04 | 0.1 | 1 | 1 | 2 | 1 | 0.495 |
|  | 6306 | 43.86 | 0.0 | 1 | 1 | 1 | 1 | 0.508 |
| 4.2 | 4690 | 97.09 | 0.057 | 1 | 1 | 1 | 1 | 0.382 |
|  | 5232 | 65.99 | 0.073 | 0 | 1 | 2 | 1 | 0.407 |
|  | 5055 | 97.82 | 0.040 | 1 | 1 | 1 | 1 | 0.452 |
|  | 4713 | 89.60 | 0.030 | 1 | 1 | 1 | 1 | 0.466 |
|  | 5250 | 16.22 | 0.074 | 1 | 1 | 2 | 1 | 0.510 |
| 4.3 | 9775 | 37.75 | 0.0 | 0.0 | 1 | 2 | 1 | 0.0103 |
|  | 9826 | 50.0 | 0.1 | 1 | 1 | 1 | 1 | 0.019 |
|  | 9775 | 50.0 | 0.0 | 0 | 1 | 2 | 1 | 0.036 |
|  | 6951 | 32.44 | 0.075 | 1 | 1 | 1 | 1 | 0.074 |
|  | 10000 | 37.75 | 0.0 | 1 | 1 | 2 | 0 | 0.075 |
| 4.4 | 1189 | 12.46 | 0.013 | 1 | 1 | 1 | 2 | 0.00080 |
|  | 3940 | 26.37 | 0.074 | 0 | 1 | 1 | 2 | 0.0017 |
|  | 8069 | 31.71 | 0.099 | 0 | 1 | 1 | 1 | 0.0019 |
|  | 3940 | 44.77 | 0.0083 | 0 | 1 | 1 | 2 | 0.0023 |
|  | 3940 | 50.0 | 0.078 | 0 | 1 | 1 | 2 | 0.0025 |

**Fig. 2.**
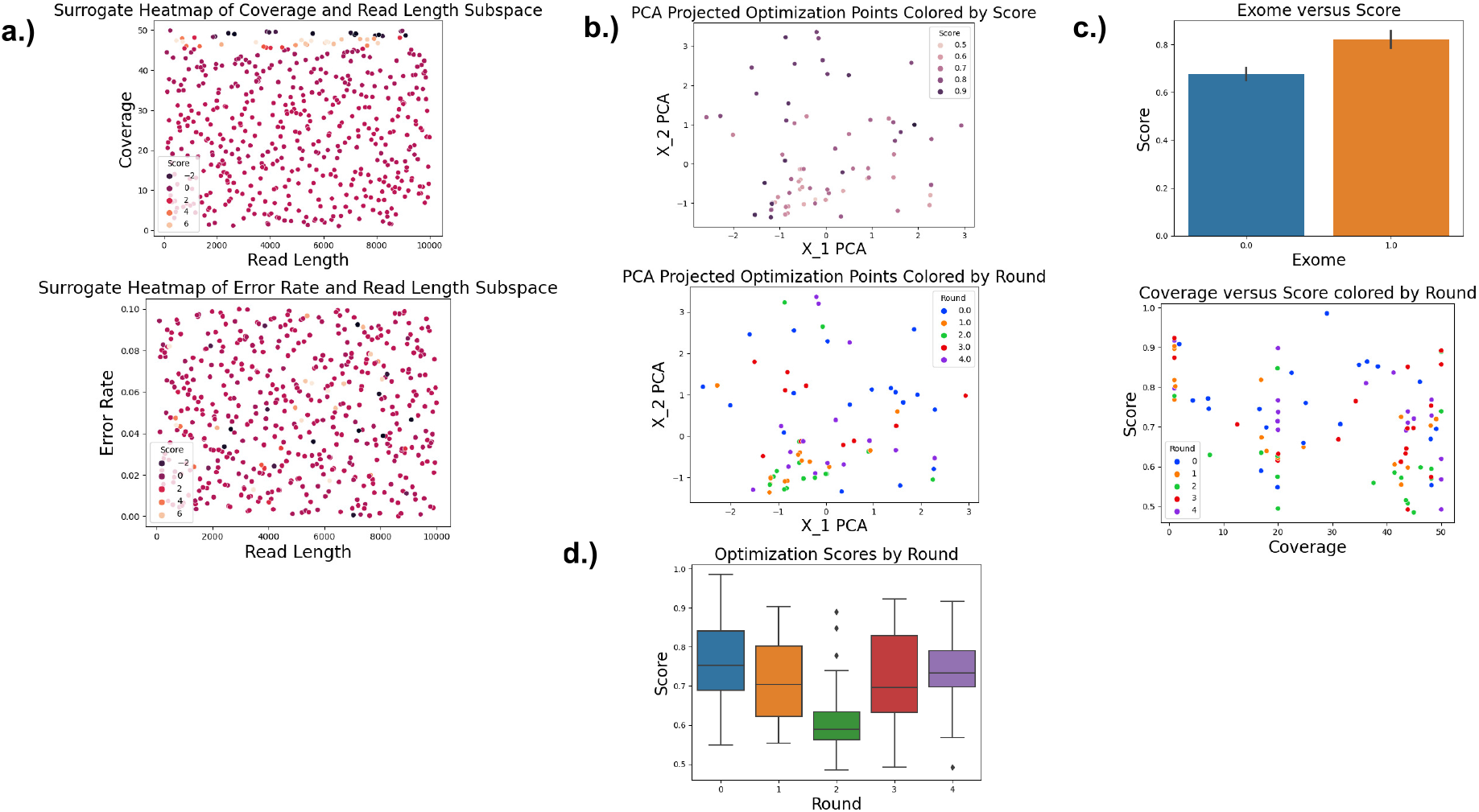
Visualizations for Experiment 4.1, an inquiry into mutation calling accuracy without a cost function. (a) 2-dimensional surrogate cross section of varying parameters, showing that high coverage values tended to be favored as expected, with read length showing a more ambiguous trend. (b) PCA plot of the points explored by the optimization algorithm. We can see that certain regions of space had better scores than others, and that the points we explored varied by the round of optimization. (c) two single parameter plots, namely exome sequencing and coverage. Exome sequencing generally had a higher score than genome, as expected; increasing coverage generally had a lower score as expected. We also see that higher coverages are explored more in later rounds than earlier rounds. (d) optimization scores by round, which showed a generally negative trend at the minimal value, but ambiguous trend when considering mean.

### 4.1 Designing a Study for Maximal Mutation Discovery

We first assume that we are presented with a tumor and wish to understand its mutational spectrum, i.e., its single nucleotide variations (SNVs) and structural variations (SVs). We assume that we have an effectively infinite experimental budget and that our cost for any study design is thus 0. Resources are encoded here in hard constraints on upper limits, rather than in the objective function. We define the loss function as:

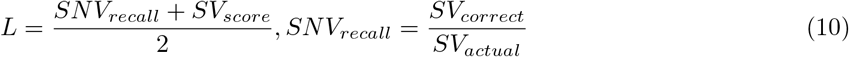

Calling the SV output locations for chromosome *i, C*_*i*_ = {(*a, b*)}, and calling our ground truth set of SV locations, *D*_*i*_ = {(*c, d*)}. The SV scoring function is then defined as:

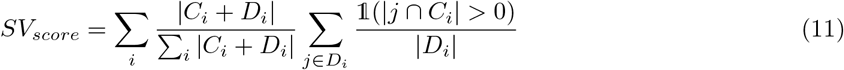

The SV portion of the loss function measures the fraction of called structural variants that overlap with the ground truth variant set. The overall loss function is intended to model the accuracy with which we can correctly detect both structural and point changes in the genome.

The optimizer’s results for this query can be viewed in Table 2. Overall, the best scores tended to be near the upper bound of the coverage parameter and to correspond to the genome sequenced with two samples with additional single-cell draws. Error Rate seemed to have a negligible effect, which makes sense as we were primarily testing for recall, and read length seemed to have a negative effect on scores near the lower extremities of the search region. Visualizations of the Exome and Coverage parameters, which appeared to be particularly significant, can be seen in Fig 2. Overall the results of the optimization procedure seemed mostly to match intuition in this experiment. With an unconstrained and broad-set query, we expect the upper-bounds of most information-containing parameters to be selected.

A univariate analysis of the study design space is likely to miss key subspaces that contain high (and low) performing designs. We see clear trends in certain 2-dimensional subspaces of parameter combinations in the surrogate function in Fig 2a.; the error rate and read length combination appears to yield mostly average performing designs with valleys and peaks interspersed. The coverage and read length combination, on the other hand, appears to show clear areas of high and low performance along the coverage axis. Similarly, a PCA projection of the generated points (Fig 2b) showed some structure, with higher score points sequestered to the top and left and the lower scoring points to the bottom and right. This may indicate some combinations of study design variables are particularly effective or ineffective. A more detailed analysis of further *n*-dimensional subpsaces or hypercubes might find subregions of higher performance.

We can also view the action of the optimization algorithm in exploring the design space. Figs 2b and 2c show that initial exploration of the entire space is fairly evenly spaced, as expected, with later rounds more concentrated in certain regions. When viewing the coverage plot in Fig 2c, we see that the initial exploration of the parameter is fairly even, but by the final round most of the samples occur towards the upper end of the permissible range. We also see that in later rounds, some “under-explored” regions of the space are still being searched due to our high exploration parameter. The optimization scores by round fluctuated considerably, with a significant number of outlier points while both the median and lowest points significantly varied. This might be due to noise from the underlying informatics pipeline, biological stochasticity, the high variance sampling strategy, or the relatively low number of points generated per round.

### 4.2 Imposing Cost Constraints on the Design Space

We now make the perhaps more tenable assumption that a researcher must consider trade offs between study efficacy and resource cost. Again, we wish to understand the overall mutational spectrum of the somatic cell population. This time, however, we are constrained by a total budget per experiment that cannot be exceeded. We also would prefer to use cheaper study designs that recover sufficient signal over designs with higher cost but equivalent efficacy. To model this situation, we create a cost function that linearly and multiplicatively scales with sequencing parameters corresponding to more depth and accuracy. This function is intended to theoretically model how costs might scale with different biological parameters. We change the overall score to incorporate a lambda penalty for increased cost:

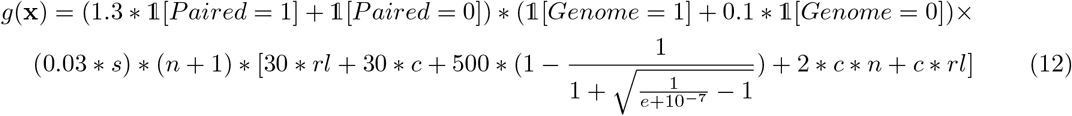

The loss function for this study is identical to that defined in Eq 10. The study design constraints in this experiment were slightly expanded to allow for increased coverage up to 100x depth, and we again replicated independent tumors arising from a stochastic process consisting of independent draws with 5 subclones and standard mutation rates consistent with tumor literature.

The lowest scoring results are shown in Table 2. As in Query 4.1, the coverage and genome parameters are near their upper bounds. Unlike the prior query, however, we see that excessive usage of certain parameters is penalized and not optimal. For example, in Fig 3a, excessively high read lengths receive high scores and the optimal read length scores tended to be within the middle of the parameter range. The addition of single cell samples did lower the scores (see Fig 3c), but some high performing samples also did not use single cell sequencing. Similarly, although we would naturally expect multiple-sample study designs to perform best, we saw that single sample study designs also performed well with respect to our overall score. This is possibly because of the multiplicative cost trade-off associated with the additional sample. We might expect to see lower parameter values with an increased cost function prioritization and for different parameter combinations to be prioritized even with a repeated run of the optimization algorithm.

**Fig. 3.**
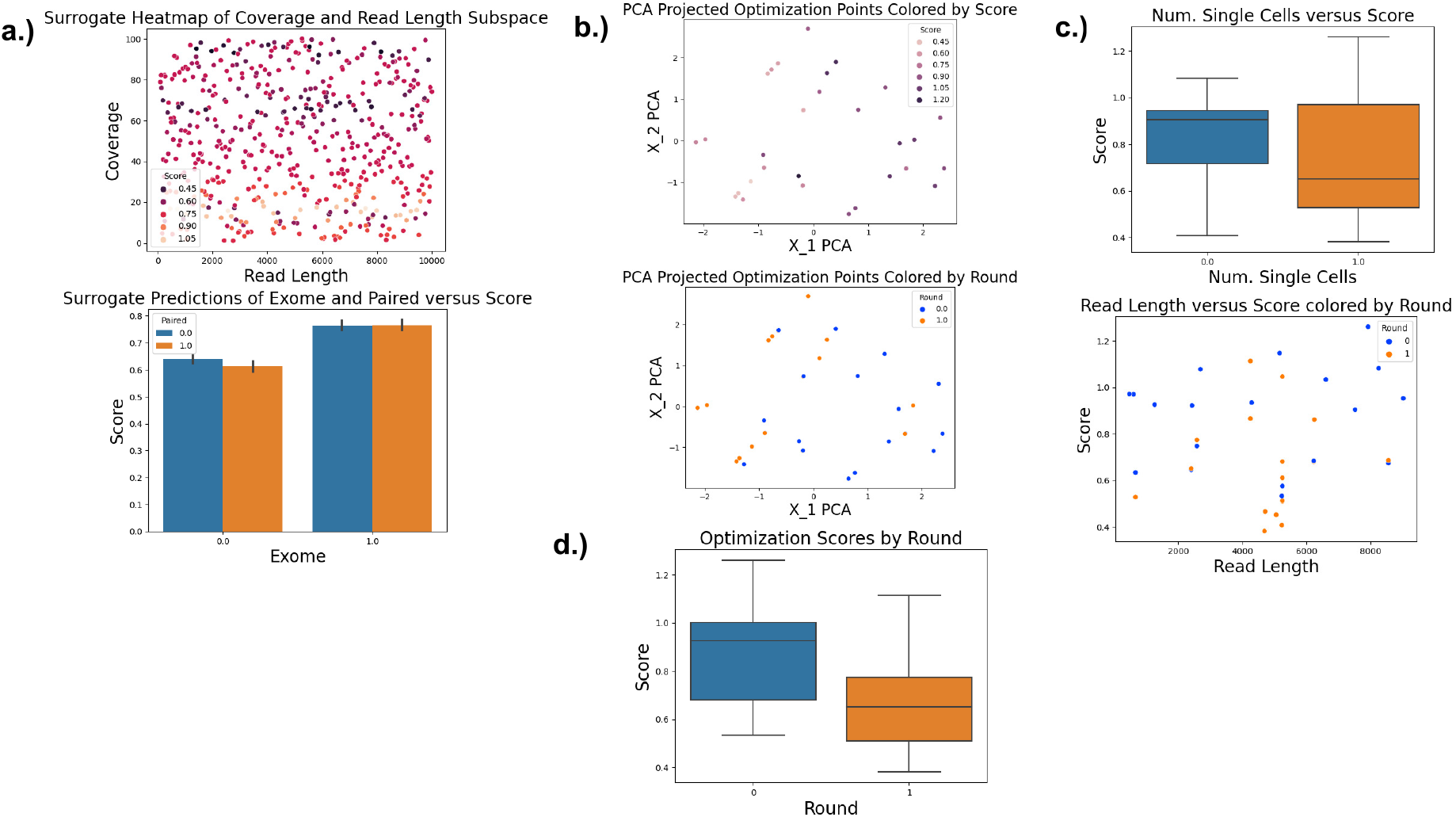
Visualizations are depicted below for Experiment 4.2, a study design problem combining a cost function with mutational calling accuracy. (a) 2-dimensional parameter trends in coverage and read length as well as exome and paired tuples of surrogate points. (b) PCA projections of the optimization points. Again, we can see certain regions of the space contain different score levels and are explored in differing fashion sequentially by the algorithm. (c) single parameter trends, where increasing single cells lowered score and read length had no clear trend. We see that read lengths in the middle of our range are explored more during the second round of optimization and produced lower scores. (d) depicts that the optimization score lowered in both mean and minimum over the course of the algorithm.

We can also see the surrogate has a different representation with the imposition of the cost function. A surrogate plot (Fig 3a) of coverage and readlength shows a more oscillatory structure with respect to coverage. Similarly, we can measure the effect of various categorical parameters. The exome parameter appears to drive down the score but the paired parameter has a much more negligible effect. We also can view the action of the optimization algorithm in terms of the points explored. Though there were only two rounds of low sample optimization, we see that the points explored in Fig 3b appear to be in different parts of the space depending on the round. With far more samples/rounds of sampling, we might expect that alternative low cost subspaces with relatively low losses might be found. It also appears that there is a fair amount of local structure in the points, as Fig 3b shows that nearby points tended to share the same scores. The true optimization function might be smoother, since the cost we assigned was deterministic and smooth and might have helped reduce noise created by the informatics calling pipeline. Similarly, the optimization scores uniformly decreased in both median and the lower percentiles (Fig 3d).

### 4.3 Measuring Structural Variation Rate

One key hypothesis in understanding how somatic mutability influences cancer risk is the idea of hypermutability processes. The aging and developing human body is constantly evolving at the genetic and epigenetic levels at some basal rates Li *et al*. (2021); Olafsson and Anderson (2021). In the case of cancers, environmental stresses or damage to key genetic regulators can cause an aberrant somatic evolutionary process that accumulate variations in characteristic fashion (Alexandrov *et al*., 2020). The most recent common ancestor of a tumor population can potentially span back many decades, implying many rounds of selection for clones generated via a hypermutability process (Körber *et al*., 2023). Given the importance of detecting somatic evolutionary processes, we seek to find a study design that will optimally approximate the rate of SV accumulation in a tumor. To measure this quantity, we use a loss function defined by the percentage difference between our called and actual SV counts.

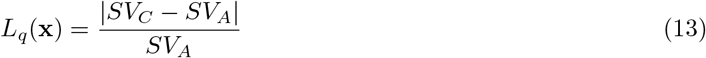

We assume that cost is fixed at 0. We apply similar biological parameters and study design parameter ranges as in the prior experiments.

The results shown in Table 2 demonstrate that the highest performing designs for this task generally maximize read length. Coverage also had a positive effect but did not see nearly the same correlation as read length. The surrogate projections (see Fig 4a) demonstrate strong negative correlation with increased read length. These results align with the commonly held perception that higher read lengths are critical to the accurate discovery of novel SVs. Values like error rate did not have a consistent effect on overall score, and the optimizer tended to oversample both the low and high extremes for error rate (4c). There were a number of outlier points that had extremely high mutational count differences. This raises the question of whether these points should be considered valid for the optimization or rerun/discounted. The sampling of points from round to round appeared to be in differing regions of the space (with the exception of the initial Latin hypercube round). The scores for the optimization appeared to trend downwards with each round, but were still rather noisy, which might again be due to the small sample size per round.

**Fig. 4.**
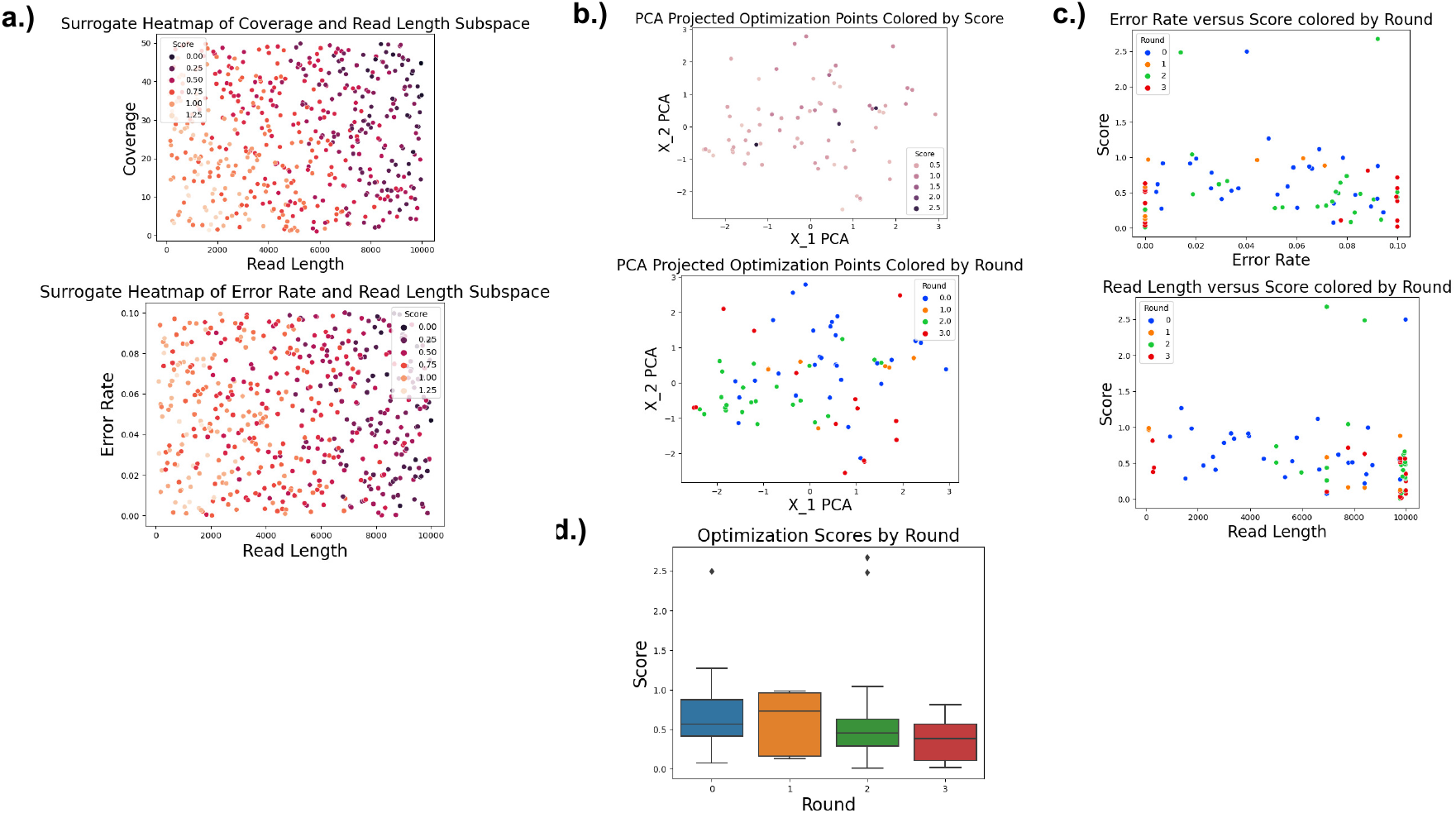
Visualizations are depicted for Experiment 4.3, a mutation rate recovery query with no associated cost function. (a) 2-dimensional parameter trends in coverage and read length as well as error rate and read length of the surrogate function; we can see a strong correlation along the read length axis. (b) optimization points explored by the algorithm projected in a PCA space, with colors depicting score and round respectively. (c) single parameter trends in error rate and read length colored by the round the point was explored. (d) shows a decrease in overall score by both mean and minimum by optimization round.

### 4.4 Generating a Minimum Viable Design for a Hypermutability Query

A common point of interest in the study of cell populations is the presence or absence of a certain feature characteristic to a particular neoplasia. For instance, one might be interested in the frequency of an SV class being in a certain range, the proliferation of a particular driver gene, or a measure of tumor purity. In each of these cases, the researcher aims to test for a specific property and would want to do so at minimal study design cost. More specifically, we now assume during the initial biopsy and sequencing of a tumor, we find that the neoplasia contains an exceptionally high rate of SVs (deletions, inversion, chromosomal segment rearrangements, etc.). We have now performed surgery and want to check for evidence of a potential relapse; to do so we want to design a screen to check for SV hypermutability at minimal cost. The query we therefore wish to answer is what is the minimal cost study design that will accurately estimate SV count/rate. We use a similar cost function as in Experiment 4.2, with our loss function this time a hinge loss — a loss of 0 is returned if the study design recovers the structural variation count with an error of up to 65 percent, otherwise 2 is returned. The primary goal with this loss function is to weight the overall score towards the cost, but only if it fulfils the desired criterion, thus generating a “minimum viable design”. Study design parameter ranges and the biological simulations are kept within standard ranges. The lost and cost functions are defined below as follows, where *SV*_*C*_ and *SV*_*A*_ denote the called and actual number of structural variants, and the study design parameters are defined in Table 1.

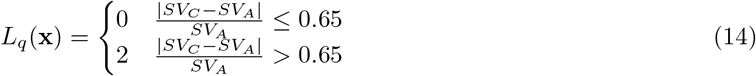

The best scores, as shown in Table 2, were highly dissimilar and appeared from distinct combinations of parameter values. Exploration of our surrogate function (Fig 5d) showed various peaks, valleys, and regions with differing score levels. Many regions of the search space and uniparameter values appeared highly dissimilar, even for geometrically close points. This potentially implies either very high noise in the results or a difficult query task. The hinge criterion may have been too strict for the level of biological noise produced in the replicates, creating a noisy ground truth scoring function that is difficult to approximate. It may also be the case that the “ground truth” scoring function for this query is complex, and to estimate such a complex query function, many more samples/rounds of sampling may be necessary. While the algorithm did explore different locations in rounds 1 and 2 (see Fig 5c,d), neither appeared to be significantly different in score, perhaps due to the strict 0 versus 2 delineation of the loss function. The results of this optimization suggest that a researcher could use the optimally produced samples in conjunction with the surrogate space as starting point to test statistical power over replicated samples. This experiment further demonstrates how the difficulty and structure in the query itself can affect optimization solutions.

**Fig. 5.**
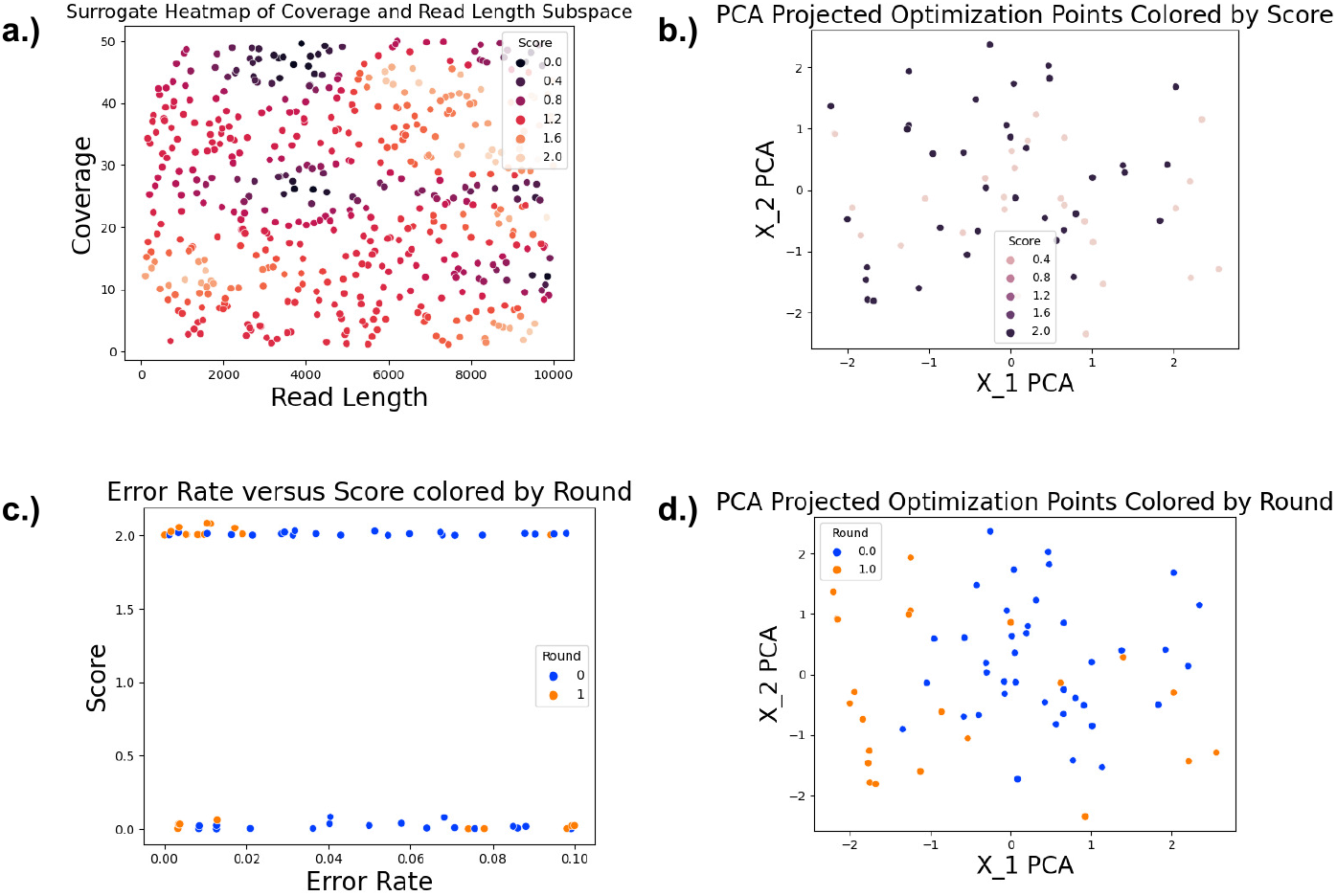
Visualizations are shown below for Experiment 4.4, a query designed to detect hypermutable cancer samples at minimal cost. Figure 4a depicts a two dimensional parameter projection of the trained surrogate points and shows a highly variable functional structure. Figure 4b similarly depicts that the explored study design points are highly dissimilar in the PCA projected space. Figure 4c shows a single parameter plot of error rate colored by round. Figure 4d depicts that the algorithm explored different parts of the space from round 1 to round 2.

## 5 Discussion

This paper seeks to address a growing problem in genomics: how to make effective use of an ever-expanding repetoire of biotechnological and computational tools so as to balance optimally our ability to discover relevant biology against resource costs. In the research world, this is a fundamental question of how to deploy finite resources most effectively. We anticipate it becoming a growing concern as well for the practice of genomic medicine, where future cancer treatment will likely involve complex genetic diagnostic and therapeutic strategies, in turn requiring design optimization strategies if they are to be used effectively. We developed a general framework for such questions and demonstrated its versatility in a series of simulated case studies, finding effective solutions that aligned with biological intuition in relatively few samples.

The present methods and use cases represent a first pass at developing a framework for posing and solving such questions, which might be further developed in many ways. We might expand these queries to complex multi-objective loss functions under complex cost constraints. The optimization procedure and solutions might be also tested through ablation studies with modifications of the lambda cost parameter, the exploration parameter, the cost function, the loss function, and budget parameters. We saw considerable noise in some of the study design samples generated, making issues of statistical power also an important area of research. Under certain conditions, provable guarantees or confidence intervals might be shown for the optimal solutions. Another key finding was that the difficulty of identifying an optimization solution is linked to the complexity of the biological space and the difficulty of the query itself. We saw that some query functions might be exceptionally smooth, whereas others might be oscillatory or even discontinuous. The importance of the query to difficulty of the optimization problem also suggests the potential of considering a parallel problem of how to design queries, which might be an optimization problem in its own right, a matter of education and guidance for users who might lack a sophisticated understanding of optimization, or perhaps a human-computer interaction problem involving a human expert working with an artificial intelligence (AI) to pose their problem most effectively to allow for its effective solution.

The broader use of optimization algorithms in biotechnology study design is relatively unexplored, with numerous opportunities for future work. Estimation of the loss function and expectation over the biological domain depends on appropriate data collection and sharing strategies. Information sharing – the idea that certain sets of data might be used as informative priors in the optimization problem and the selection of new data – could be of promise. The structure of the surrogate function is an important consideration in both the optimization procedure as well as query analysis. Ideas from semi-supervised or federated learning — where certain sets of data are created through a combination of labeled, un-labeled, and synthetic data from various sources (Zhang *et al*., 2021; Wang *et al*., 2013) — may be of aid. Other algorithmic strategies, such as Monte Carlo or variational inference to compute the expectation in equation (1), may also be of use. Active learning — adapting a study design optimally as the study proceeds — might also prove effetive. Experimental and clinical validation studies would provide further support for the algorithm, for example, in continually optimizing study designs over the clinical course of a tumor’s progression.

## Acknowledgements

Research reported in this publication was supported by the National Human Genome Research Institute of the National Institutes of Health under award number R01HG010589. The content is solely the responsibility of the authors and does not necessarily represent the official views of the National Institutes of Health.

## A1 Appendix

### A1.1 Programming Implementation and Usage Guide

All code used for the optimization program can be found at: github.com/CMUSchwartzLab/StudyDesignOptimization. The optimization code is stand-alone in that it can be easily run provided the simulation code is replaced. All required packages can be installed in a conda environment, and the parameters (e.g. budget, cost function, biological parameters, etc.) of the optimization should be changed to best suit the user’s query. The full optimization-simulation workflow is tailored to our compute system; as such, efficient replication of the workflow would require changes to the computer code to best suit the user’s compute cluster. To fully generate automated results through simulation, a number of genome aligners and variant callers need to be installed as well as modifications to the paralellization scheme and storage containers for reference and output genomes. The simulation of clonal genomes and scientific biotechnology was generated from a lightly modified version of a clonal evolution simulator presented in (Srivatsa *et al*., 2023). The major aligners used in the experimental section were BOWTIE2 (Langmead *et al*., 2009), bwa-mem (Li, 2013), and minimap (Li, 2018). The variant callers used were strelka (Saunders *et al*., 2012) and dysgu (Cleal and Baird, 2022).

### A1.2 Algorithm Pseudocode and Informational Parameter Tables

#### Algorithm 1: Complete Optimization Algorithm Pseudocode

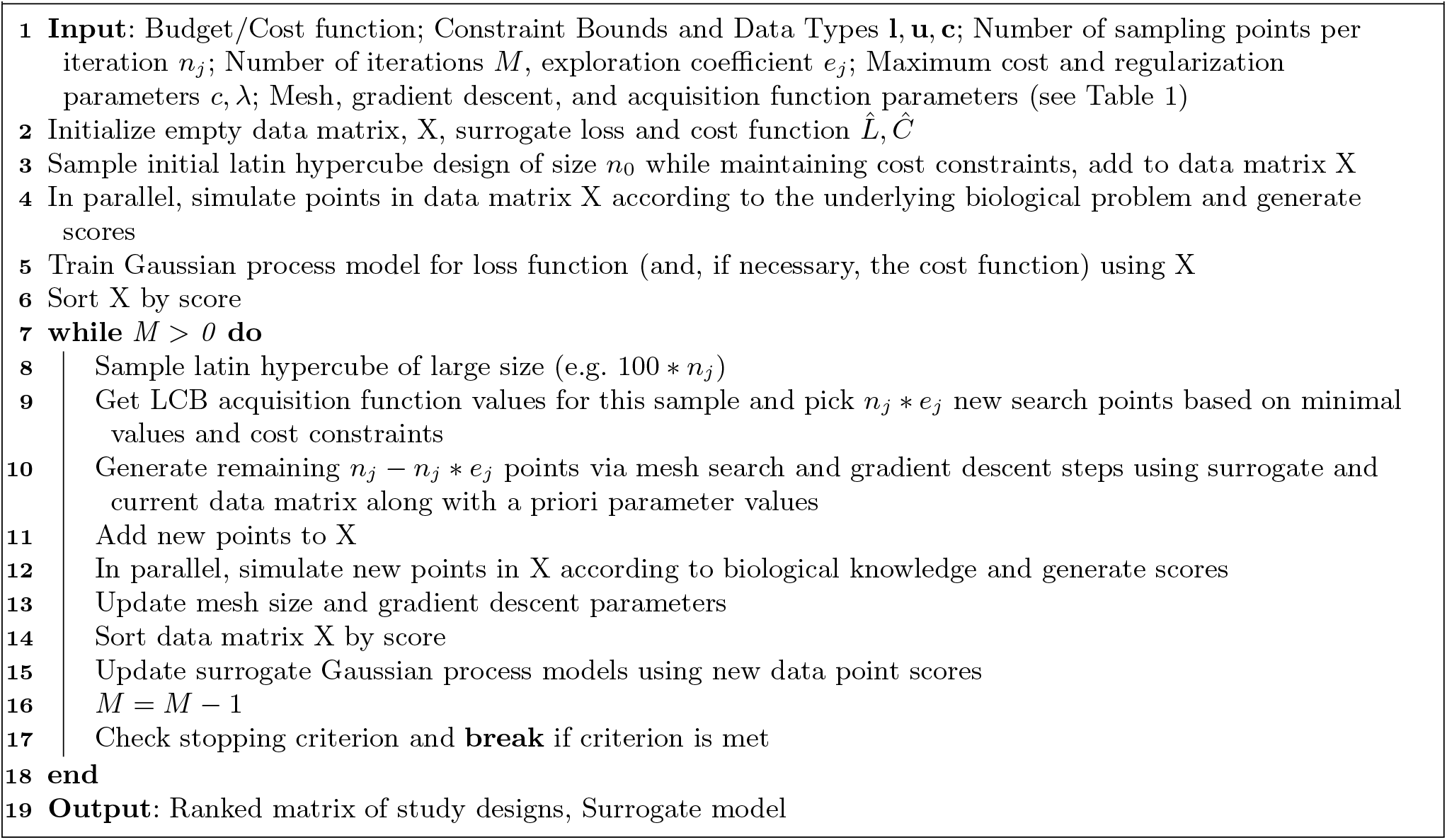

**Table A1.** A summary of the main algorithm parameters is depicted in the table above. Parameters like the bounds and cost function indicate user inputs critical to the optimization problem. Technical parameters like the gradient descent step and mesh size impact the action of the algorithm.

| Parameters | Symbols | Description |
| --- | --- | --- |
| Lowerbounds, Upperbounds, Categorical Flags | $\mathbf{l}, \mathbf{u}, \mathbf{c}$ | Hard constraints on the range of each parameter and whether they take discrete or continuous values |
| Experimental Budget, Number of Samples per Iteration | $M, n_j$ | The total experimental budget defines the number of iterations of the algorithm we can perform (i.e. the number of sets of data we can generate) and the number of samples per iteration dictates the total number of individual study designs we can generate per round. |
| Gradient Descent Step, Perturbation Vector, Scaling Factor, Number of Gradient Descent Sample Points | $\alpha_g^j, \mathbf{e}_j, \gamma, n_g^j$ | These parameters define the numerical estimation of the gradient as well as the size of the gradient iteration step. The perturbation vector and scaling factor are used to compute a finite difference equation for the gradient at a certain point, and a new point is generated from the current point and gradient descent step. The number of gradient descent points dictates the number of current minimal points whose gradients are estimated for the next iteration. |
| Exploration Coefficient, Acquisition Variance Parameter | $e_j, \alpha_v^j$ | The exploration parameter assigns a fraction of points to explore via our acquisition function derived from the Gaussian process as opposed to local search of already promising points. The acquisition variance parameter emphasizes the exploration of high variance points at large values. |
| Mesh Size, Directional Polling Set, Discrete Neighborhood | $\Delta_j, D_j, \mathcal{N}_D$ | The mesh size assigns a range for the lattice search, while the directional polling set and discrete neighborhood definitions assign how we explore local points from the mesh center. |
| Cost Function, Loss Function | $g(x), L(x)$ | The cost function defines the cost for a single study design and can either be given in closed-form to the optimization program or approximated by a surrogate function from data. The loss function describes the efficacy of a study design and must be computed through experiment or simulation. |
| Maximum Cost, Lambda Regularization Parameter | $c, \lambda$ | The maximum cost parameter describes the limit in cost of a study design and implicitly assigns a feasible region to our study design space. The lambda regularization parameter balances a trade off between cost and loss in our overall optimization output (i.e., how much emphasis we should place on gains in performance versus increases in cost). |

**Table A2.** Biological Parameters that influence the instance of somatic evolution are written and described in the table above.

| Parameter | Symbol | Units | Description |
| --- | --- | --- | --- |
| Effective Population Size | $N_e$ | Cells | Total cellular population of the region to be sampled. Impacts coalescent times |
| Number of Clones | $k$ | Clones | Number of distinct genetic somatic cell populations to be sampled |
| Mutation Rate Lists | $M_i$ | Mutation events per locus per cell division | Lists for each variant class, defining rates per locus per cell division |
| Mutation Size and Location Distributions | $S, z(x)$ | Number of Bases, None | Each mutation type can be tuned over size distributions and single base substitutions can be tuned over signature distributions |
| Number of Tumors | $t$ | Tumors | Number of distinct sites of somatic evolution to be sampled |
| Dirichlet Concentration, Clonal Frequency Distribution | $\alpha, g(k)$ | None, None | Parameter of a Dirichlet process which is used to derive the concentration of the baseline distribution in a sample. A high value will lead to approximately uniform sampling of clones during sequencing, whereas a low value would favor very uneven clonal frequencies. The clonal frequency distribution is the baseline distribution at high $\alpha$ |

**Table A3.** Study Design parameters influencing the construction of a research study and associated genetic toolbox are written and described in the table above.

| Parameter | Symbol | Data Type | Description |
| --- | --- | --- | --- |
| Number of Samples | $s$ | Samples | Number of distinct tissue biopsies to be drawn from each “tumor” site |
| Type of Sample | $sample$ | None | Defines how a sample is taken from the cell population (generally solid or liquid biopsy, though other methods like liquid capture microdissection are less commonly used) |
| Number of sequencing protocols taken on the $i^{th}$ sample | $l_i$ | Integer | Number of individual sequencing protocols taken on the sample $i$ |
| Number of single cells used in a sequencing protocol | $n$ | Integer | Number of single cells operated upon under a singular sequencing protocol. If zero, no single cells are generated and only bulk sequencing is performed. |
| Read Length | $rl$ | Integer – number of Bases | Size of reads to be generated from sequencer |
| Fragment Length | $fl$ | Integer – number of bases | Defines a superstring from which reads are derived during the sequencing process |
| Depth/Coverage | $c$ | Integer – reads per Base | Average number of times each nucleotide of the genome is sequenced |
| Error Rate | $e$ | Float – Fraction of Incorrectly Sequenced Bases | Rate of Incorrectly sequenced nucleotides |
| Paired-End/Single-end | $Paired$ | Boolean | Binary parameter describing whether the reads are paired end or single end. Paired-end reads have two related reads derived from the same fragment. |
| Whole Genome/Whole Exome/Targeted Sequencing | $Genome$ | Boolean | Binary parameter describing whether the sequencer extracts genes from the entire genome or only a subset of the genome |
| Informatics Tools | $Informatics$ | Categorical/numerical | A parameter describing the type of informatics software used to analyze a sequencing protocol for information. Notably the encoding is flexible, and could be represented many different ways. |

